# Glycolytic inhibitor 2-Deoxy-D-glucose attenuates SARS-CoV-2 multiplication in host cells and weakens the infective potential of progeny virions

**DOI:** 10.1101/2021.06.12.448175

**Authors:** Anant Narayan Bhatt, Abhishek Kumar, Yogesh Rai, Neeraj Kumari, Dhiviya Vedagiri, Krishnan H. Harshan, Vijayakumar Chinnadurai, Sudhir Chandna

**Affiliations:** Institute of Nuclear Medicine & Allied Sciences, Delhi, India.; CSIR-Centre for Cellular and Molecular Biology, Hyderabad, India 500007; Academy for Scientific and Innovative Research (AcSIR), Ghaziabad-201002, India

**Keywords:** SARS-CoV-2, Glycolysis, 2-DG, Unglycosylation, COVID-19

## Abstract

The COVID-19 pandemic is an ongoing public health emergency of international concern. While a lot of efforts are being invested in vaccinating the population, there is also an emergent requirement to find potential therapeutics to effectively counter this fast mutating SARS-CoV-2 virus-induced pathogenicity. Virus-infected host cells switch their metabolism to a more glycolytic phenotype. This switch induced by the virus is needed for faster production of ATP and higher levels of anabolic intermediates, required for new virion synthesis and packaging. In this study, we used 2-Deoxy-D-glucose (2-DG) to target and inhibit the metabolic reprogramming induced by SARS-CoV-2 infection. Our results showed that virus infection induces glucose influx and glycolysis resulting in selective high accumulation of the fluorescent glucose/2-DG analogue, 2-NBDG in these cells. Subsequently, 2-DG inhibits glycolysis in infected cells thereby reducing the virus multiplication and alleviates the cells from virus induced cytopathic effect (CPE) and cell death. Herein, we demonstrate that the crucial Nglycosites (N331 and N343) of RBD in spike protein of progeny virions produced from 2-DG treated cells were found unglycosylated and defective with compromised infectivity potential. In line with earlier reported observations, our study also showed that 2-DG mediated metabolic inhibiton can attenuate SARS-COV-2 multiplication. In addition, mechanistic study revealed that the inhibition of SARS-COV-2 multiplication is attributed to 2-DG induced un-glycosylation of spike protein. Our findings strengthen the notion that 2-DG effectively inhibits SARS-CoV-2 multiplication. Therefore, based on its previous human trials in different types of Cancer and Herpes patients, it could be a potential molecule to study in COVID-19 patients.

## Introduction

The coronavirus pandemic is an ongoing global outbreak of COVID-19 disease caused by severe acute respiratory syndrome coronavirus 2 (SARS-CoV-2) and recognized as Public Health Emergency of International Concern by the World Health Organization (WHO). The highly infectious virus pathogenesis caused by SARS-CoV-2 is a respiratory disease occurring in three Phases 1) Virus multiplication, 2) Immune hyper-reactivity and 3) Pulmonary destruction [1]. The faster multiplication of COVID-19 virus-driven damage to epithelial cells results in pulmonary destruction and cytokine storm [2]. An immense effort has already been made and many are underway worldwide in the direction of developing effective anti-virus therapeutic against this fast mutating virus to control the disease severity and mortality [3]. Currently, newly identified therapeutics are under investigation and there is no established therapy for the treatment of COVID-19. Treatment is largely based on supportive and symptomatic care. Therefore, the development of effective therapeutic against COVID-19 is urgently needed.

Upon infection, the virus hijacks and reprograms the host cell metabolism for its own rapid multiplication as new virion assembly involves very high demand and turnover of nucleotides and lipids [4]. This enhanced demand of virus-infected host cell is solely fulfilled by induced anabolic reactions to synthesize more nucleotide and lipid using glucose and glutamine as substrate [5]. Like cancer cells, the enhanced aerobic glycolysis, popularly known as the Warburg effect was also observed in SARS-CoV-2 infected host cells, which satisfy the elevated anabolic demand [6, 7]. In addition to providing direct substrates for virion assembly, adjustments to metabolic pathways are also required to provide ATP at a rapid rate for the high energy costs of virus genome replication and packaging in host cells [5]. While oxidative phosphorylation provides significantly more ATP per glucose, glycolysis is a much faster process for providing ATP, rapidly [8]. However, utilizing glycolysis as the main metabolic pathway requires an increased influx of extracellular glucose using enhanced expression of glucose transporters viz. GLUT1, GLUT4 etc. to match the increased metabolic rate [8]. Unlike normal cells, which show more plasticity in their metabolism by relying on both glycolytic and mitochondrial pathways, SARS-CoV-2 infected cells operate predominantly on glycolysis for bioenergetic and anabolic demand due to compromised mitochondrial respiratory function [6]. Based on these facts, the intervention at the level of SARS-CoV-2 induced host cell metabolism is a promising target to exploit as a potential strategy for developing anti-virus therapy to control the progression of the disease and thereby the pandemic.

The glycolytic inhibitor, 2-Deoxy-D-glucose (2-DG) is a synthetic analogue of glucose, which blocks glycolysis at the initial stage and cause depletion of ATP, anabolic intermediates required for virus synthesis and glucose derivatives used in protein glycosylation [9]. It is demonstrated in earlier studies that inhibition of reprogrammed metabolism of virus-infected host cells using 2-DG potently impairs virus multiplication by reverting anabolic reprogramming [6,10–12]. Hence, we used 2-DG to target the selective and predominant metabolic pathway of the virus-infected host cell, glycolysis to inhibit virus multiplication.

In this study, we studied the effect of SARS-CoV-2 infection on the alterations of host cell metabolism with respect to enhanced expression of transporters and enzymes responsible for glucose uptake and glycolysis pathway. We found 2-DG treatment not only inhibited the virus multiplication but also lead to the formation of defective virions, which have reduced ability to infect the newer cells resulting in overall effective inhibition of virus growth.

## Results

### SARS-CoV-2 infection induces cellular glucose influx and glycolysis

To evaluate the effect of SARS-CoV-2 infection mediated reprogramming of the host cell metabolism towards enhanced glucose utilization, we estimated the glucose uptake using 2- NBDG, a fluorescent glucose/2-DG analogue. Using fluorescence imaging of cells, we found SARS-CoV-2 infection in Vero E6 cell leads to profound accumulation of 2-NBDG as compared to uninfected cells (Fig. 1A), which was confirmed by image analysis and quantitative estimation of 2-NBDG fluorescence intensity (Fig.1 B-C). Additionally, the enhanced glucose uptake of infected cells was confirmed by significantly increased levels of key glucose transporter proteins GLUT1, GLUT3 and GLUT4, corroborating the infection mediated upregulation of glucose influx in virus-infected cells (Fig.1 D). Besides, significantly enhanced levels of key glycolytic enzymes viz. Hexokinase-II (HK-II), Phsopho-fructokinase-I (PFK-1), Pyruvate Kinase (PKM-2) and Lactate dehydrogenase (LDH-A) were observed in virus-infected cells (Fig.1 D). Analysis of glucose uptake and glycolytic enzymes revealed that virus-infected cells accelerate the glycolysis to meet the high energy demand, supported by the significant increase in ATP production, compared to uninfected cells (Fig. 1C). Moreover, the higher protein levels of glycolytic enzymes are responsible for the regulation of diverting the glucose through glycolysis and conversion of pyruvate to lactate after glycolysis, substantiating the evidence of reprogrammed metabolism towards increased glycolysis in SARS-CoV-2 infected Vero E6 cells.

**Figure 1.**
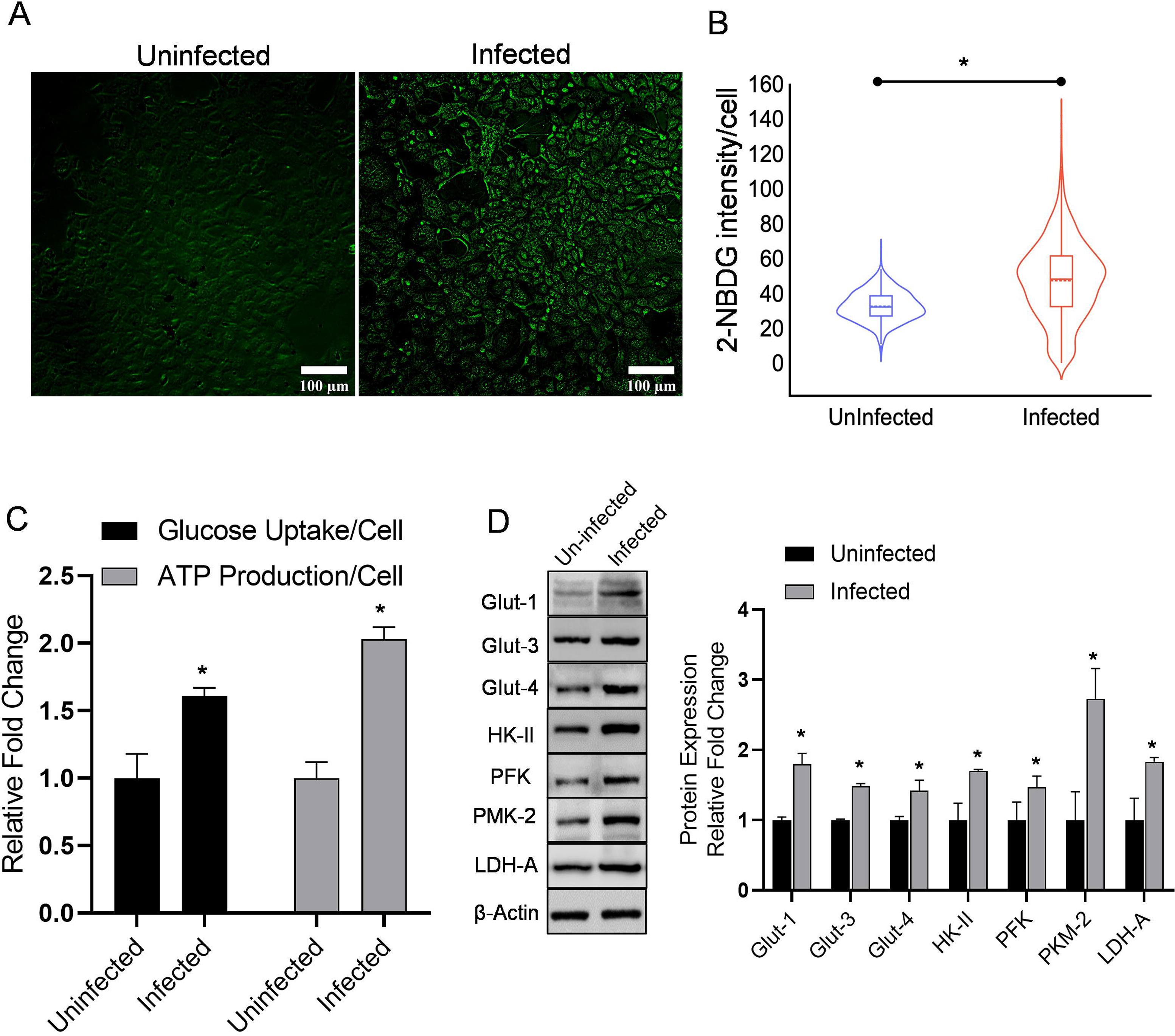
SARS-CoV-2 infection induces glycolysis in Vero E6 cells. **A.** Photomicrographs show the 2-NBDG (200µM) staining in Vero E6 cells at 48 h post-infection captured under 10×10 X magnification. **B.** An intracellular fluorescence uptake was quantified (derived information of A) using software toolbox CELLSEGM on MATLAB and the graph presented as 2-NBDG fluorescence intensity/cell. **C.** In a similar experimental condition, 2-NBDG uptake and ATP were estimated and the graph represents the relative fold change (RFC) in the indicated groups. **D**. Immunoblot and densitometric evaluation indicating the glucose influx and glycolytic profile of SARS-CoV-2 infected Vero E6 cells and presented as RFC with respective control. Data represent the mean ± SD of at least two independent experiments performed in triplicates. *p*<*0.05 presented with respect to uninfected control (student’s *t*-tests paired analysis).

### SARS-CoV-2 infection mediated enhanced glucose uptake is selective to infected cells

To further deepen our understanding about the selectively higher glucose uptake in SARS-CoV-2 infected cells and low in uninfected or minimally infected cells, we tested the 2- NBDG uptake in the co-culture of cells. A similar cell line Vero was selected, which showed compromised infection and poor multiplication efficiency of SARS-CoV-2 virus upon infection with similar MOI (0.3) with respect to Vero E6 (Suppl. Fig. 1) [13]. To differentiate between the cells, Vero cells were stained with CTR (cell tracker red) and cocultured with Vero E6 cells, the co-culture was infected 24h later with SARS-CoV-2 (Fig. 2A). Interestingly, the basal level uptake of 2-NBDG in infected Vero cells showed a minimal increase compared to respective control visualized as merged green (2-NBDG) and red (CTR) emission of Vero cells (Fig. 2A). Whereas, the mostly acquired intracellular green fluorescence in co-culture corresponds to a significant increase of 2-NBDG uptake by Vero E6 cells following infection, compared to uninfected cells (Fig.2B). Quantitative estimation of fluorescence intensity from multiple images of these samples confirmed that 2-NBDG uptake is significantly favoured by SARS-CoV-2 infection, selectively in highly infected Vero E6 cells (Fig.2B). These findings suggest that SARS-CoV-2 infection favours increased glucose as well as a 2-DG influx in cells.

**Figure 2.**
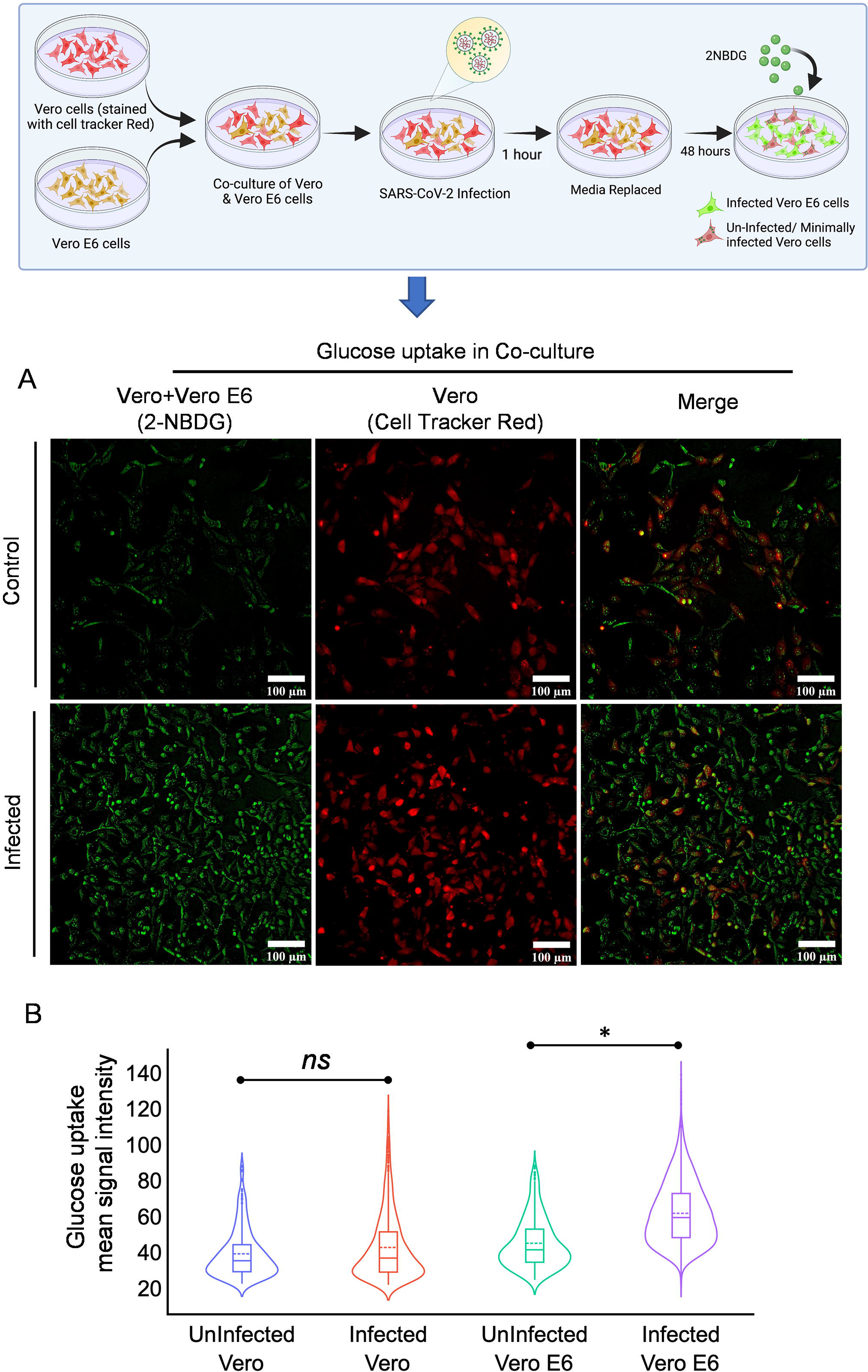
Glucose uptake is selectively more in highly infected cells. **A.** The graphical presentation depicts the methodology of co-culture of Vero and Vero E6 cells. Vero cells with relatively poor ability of infection uptake were stained with CellTracker™ Red and co-cultured with Vero E6 cells to test if 2-NBDG accumulation is preferably favoured by SARS-CoV-2 infected cells. Photomicrographs in the left panel show the 2-NBDG (green) uptake in Vero and Vero E6 cells of indicated treatment groups whereas, the middle panel are representative of CTR (Red) stained Vero cells in the same field and right panel micrographs are the merged representation of left and right panel for the better appreciation of differential 2- NBDG accumulation in Vero and Vero E6 cells. **B.** The violin plot shows the mean signal intensity of 2-NBDG in the indicated treatment groups (quantitatively derived from A). *p*<*0.05 presented with respect to uninfected control (student’s *t*-tests paired analysis).

### Effect of glucose antimetabolite 2-DG on cell proliferation and viability

Since SARS-CoV-2 infected cells predominantly showed increased glucose metabolism. Therefore, we hypothesized that selective inhibition of glucose metabolism using 2-DG may reduce the SARS-CoV-2 infection in these cells. To validate our hypothesis first we examined the safe concentration range of 2-DG in Vero E6 cells using SRB assay to understand the effect of 2-DG on cell proliferation. The decrease in cell growth was observed in 2-DG treated cells and notified as inhibition range of 1-10% at 0.1-0.5mM, 10-30% at 1-5mM and reached maximum inhibition ∼40-50% at 10-100mM (Fig.3 A) in Vero E6 cells. It was interesting to observe that 5-100 mM showed a near saturation range of growth inhibition (Fig.3 A), indicating that the observed growth inhibition is not due to the toxicity of 2-DG. The glycolysis inhibitor 2-DG is known to inhibit the proliferation of rapidly multiplying cells and exert a cytostatic effect. Therefore, we analyzed the effect of 2-DG on cell proliferation using the CFSE probe. Relatively more CFSE fluorescence in 1mM and 5 mM treated cells as compared to their respective time control indicates the slower proliferation in 2-DG treated Vero E6 cells (Fig.3 C&D). Further, to understand deeply about the 2-DG induced growth inhibition, we analyzed the reproductive potential/ clonogenicity following 2-DG treatment by performing a macrocolony assay (Fig.3 B). The non-significant change in the clonogenicity between control and 2-DG (1 and 5mM) treated cells showed that 2-DG induced growth inhibition is due to reduced proliferation and cytostatic effect and not due to cytotoxic effect. Therefore, the highest used concentrations of 2-DG (upto 5mM, in most of the experiments) was found to be safe for Vero E6 cells.

**Figure 3.**
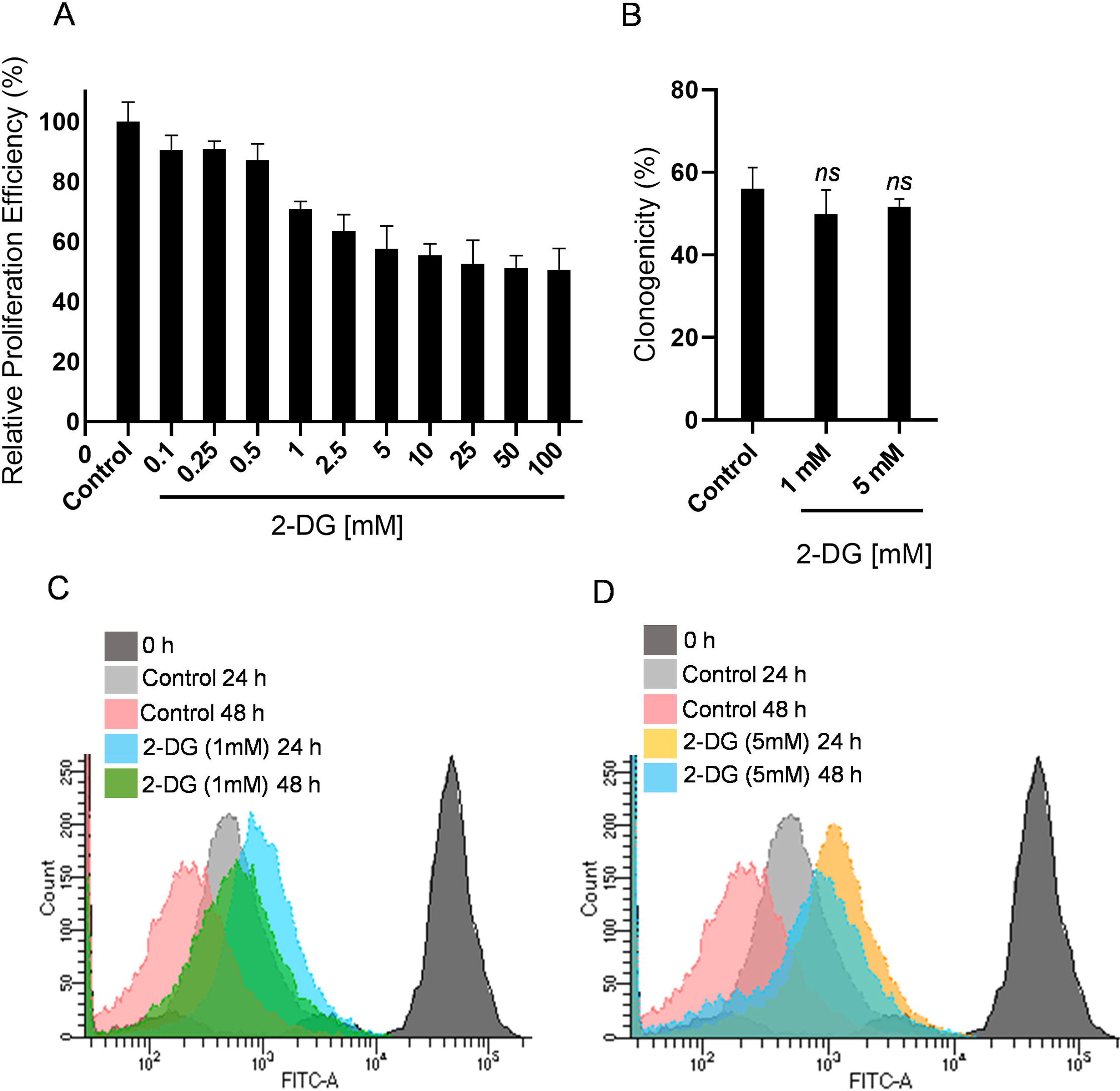
2-DG treatment reduces the proliferation efficiency of Vero E6 cells. **A.** SRB assay was performed to examine the effect of 2-DG on cell proliferation. The graph shows the relative proliferation efficiency (%) of Vero E6 cells at the different indicated concentrations of 2-DG with respect to control. **B.** The bar graph represents the reproductive potential/clonogenicity of 2-DG treated cells at 1 and 5 [mM], with respect to control. **C-D.** CFSE proliferation was performed, and a differential overlay graph obtained in the indicated treatment groups shows the proliferation dependent fluorescence shift in Vero E6 cells.

### 2-DG inhibits SARS-CoV-2 replication and ATP production

Initially, we assessed the concentration-dependent growth inhibition effect of 2-DG on SARS-CoV-2 (B.6 variant; earlier in May 2020) multiplication (GISAID ID: EPI_ISL_458075; virus ID- hCoV-19/India/TG-CCMB-O2-P1/2020) in Vero cells [14]. The value of half-maximal inhibitory concentration (IC50) on viral E and RdRp gene multiplication was estimated as 1 and 0.6 mM in the supernatant and cell-bound fractions respectively (Fig. 4A & B). Next, we examined the effect of 2-DG on virus multiplication of B.1.1 variant of SARS-CoV-2 (HV69/70 mutation in S gene, isolated from a patient sample ID INMAS/nCoV/8415) by analyzing the RdRp gene and N gene from secreted virions in cell medium (Fig. 4C) and cell-bound virions in cellular fraction (Fig. 4D). 2-DG strongly inhibited SARS-CoV-2 growth in Vero E6 cells (Fig. 4 C&D). The estimated IC_50_ value was found to be 0.75 mM and nearly 0.9 mM in released and cell-bound fractions, respectively which was nearly similar to B.6 variant. It was also observed that 2-DG reduces the level of virus proteins namely nucleocapsid (N) and spike (S) protein in a dose-dependent manner (Fig. 4E). Further, in 2-DG treated cells reduced viral load and protein expression was correlated with the significant reduction in cellular ATP content following SARS-CoV-2 infection (Fig. 4F). It is also pertinent to note here that 2-DG treatment alone did not show a considerable reduction in ATP production (Fig. 4F). These results indicate that 2-DG inhibits glycolysis and thereby faster ATP production, eventually preventing virus multiplication in host cells.

**Figure 4.**
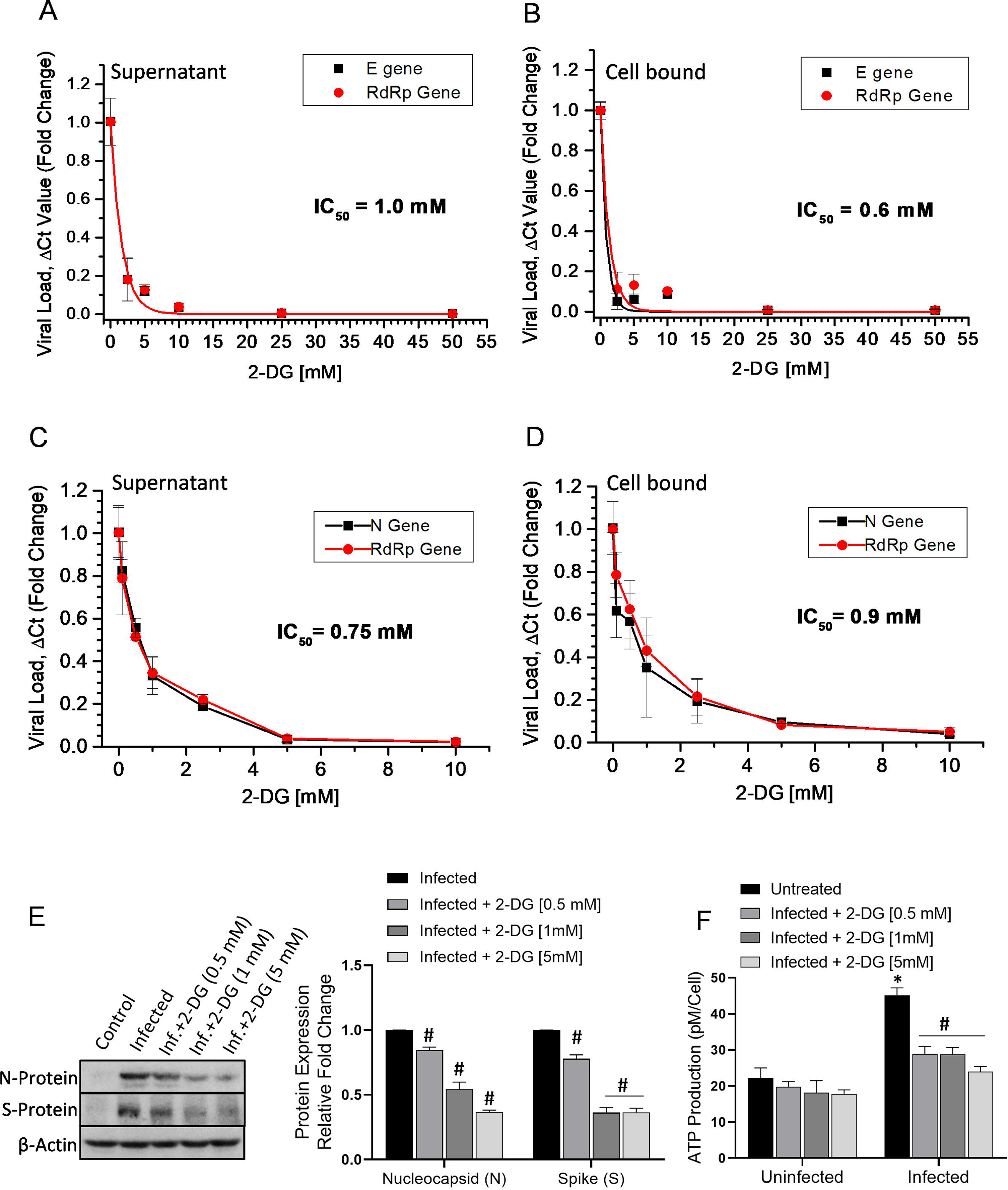
2-DG treatment inhibits SARS-CoV-2 replication in Vero and Vero E6 cells A-D. RT-PCR of viral E and RdRp genes (**A&B**) in SARS-CoV-2 infected (B.6 variant) Vero cells; N and RdRP genes **(C&D)** in infected (variant B.1.1) Vero E6 cells were performed in the supernatant of the infected culture and its cellular fraction. The concentration-dependent inhibitory effect of 2-DG is presented as relative fold change (RFC) in reduced virus load (ΔCt) with respect to the infection control group. **E.** Immunoblotting of virus N and S protein was performed with the whole cell lysate of indicated treatment groups and the densitometry graph shows the RFC in 2-DG treated [0.5, 1 & 5mM] fraction with respect to infection alone. **F.** ATP was estimated at 48 hrs post-infection in the indicated treatment groups followed by normalization with respective cell numbers and graph presented as ATP production pM/Cell. Data represent the mean ± SD of four independent experiments where each condition was analysed in triplicates. *p*<*0.05 compared to control and ^#^p<0.05 with respect to infection alone (student’s *t*-tests paired analysis).

### 2-DG inhibits SARS-CoV-2 mediated cytopathic effect and cell death

The virus infection-induced cellular deformity, cell lysis and cell death of host cells are very well characterized phenomenon and commonly known as cytopathic effect (CPE). After analyzing the effect of 2-DG on virus multiplication, next, we analyzed the effect of 2-DG on SARS-CoV-2 infection mediated cytopathic effect (CPE) and induction of cell death. Upon microscopic examination, a profound change in cell morphology was observed in infected cells at 48 hours post-infection, whereas this effect tends to be low at 1 mM and visibly absent at 5 mM 2-DG, suggesting considerable low CPE following 2-DG treatment (Fig. 5A). The SARS-CoV-2 infection-induced change in cell death index and integrity of cell membrane was further analyzed by fluorescence imaging of dual DNA binding stain acridine orange (AO: Green) and ethidium bromide (EB: Red), viewed and analyzed in terms of the rate of selective dye ingress and accumulation depending upon the physiological state of the cell. The cellular accumulation and increased emission peak of AO were facilitated predominantly by infected cells only and appeared as a bright green nucleus indicating a higher early apoptotic phase population known to be characterized as condensed and fragmented chromatin [14]. Additionally, in these cells, substantial loss to the cytoplasmic membrane integrity and more necrotic population were also visualized by a selective increase in EB stained populations. In contrast, 2-DG treated cells showed a significant reduction in the early apoptotic population and considerable restoration of membrane integrity confirmed by the quantitative evaluation of cellular fluorescence intensity of both AO/EB stained cells (Fig. 5 B-C). Together these results suggest that 2-DG treatment enable cells to overcome the cellular stress caused due to SARS-CoV-2 multiplication and thereby reduces cell death.

**Figure 5.**
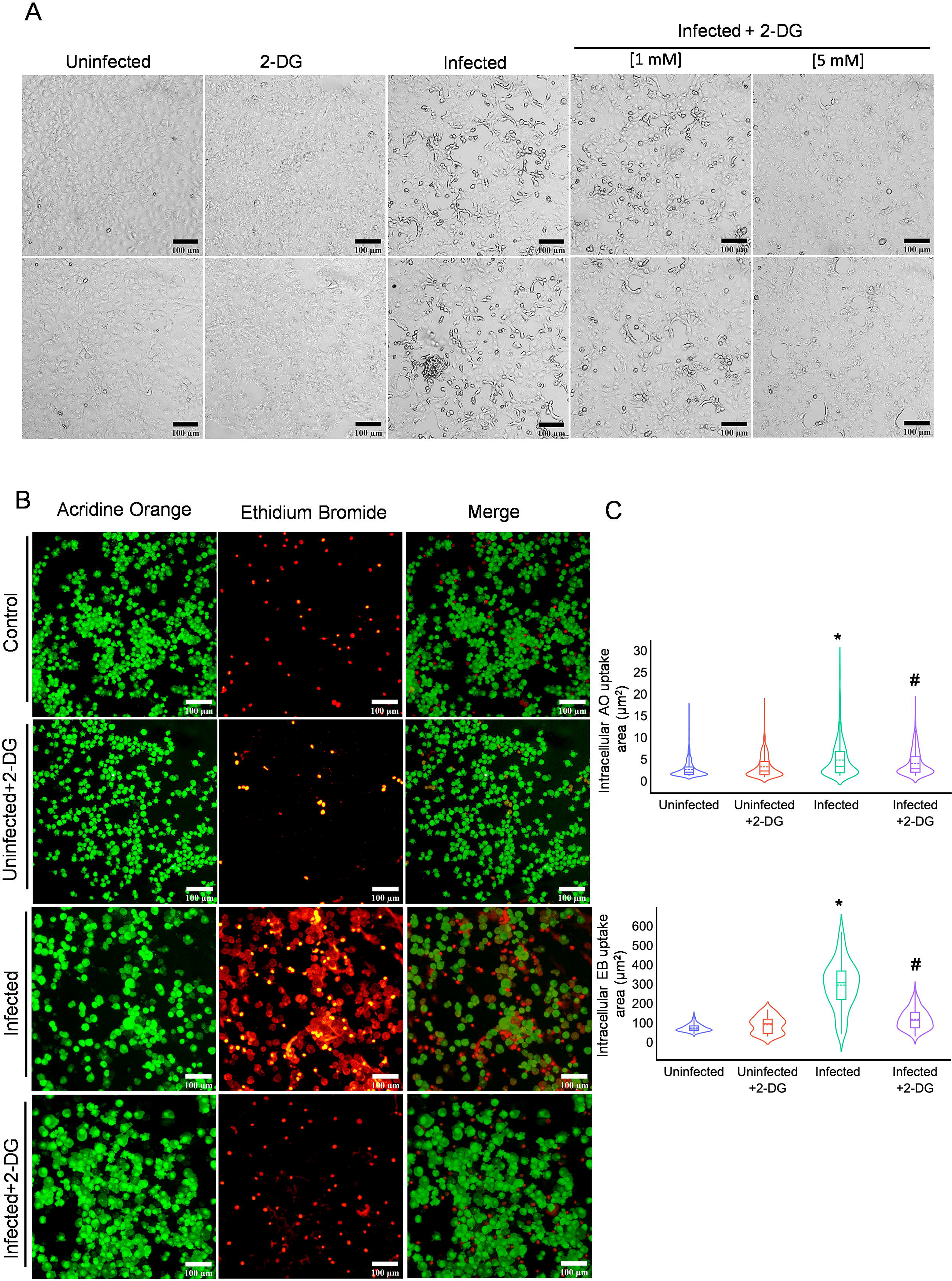
2-DG treatment reduces SARS-CoV-2 induced cell death. **A.** Photomicrographs show the differential change in cellular morphology upon virus infection in the indicated treatment groups performed at 48 h post-infection in Vero E6 cells. **B.** In the similar experimental condition fluorescence images of acridine orange (AO) and ethidium bromide (EB) (AO/EB; 100 μg/ml each; 1:1) presented using red and green emission channel respectively in Vero E6 cells (suspension form) captured under 10 × 10 X magnification. **C.** Photo-morphometric analysis (derived from B) was carried out using algorithms from CELLSEGM and Image Processing toolbox in Matlab 2020a and violin plot indicating the difference in intracellular uptake and acquired area of both AO/EB stained cells calculated from (1000 cells) obtained from different field of view in the uninfected, 2-DG treated, infected and infected +2-DG treated cells. Data represents the mean ± SD of at least two independent experiments performed in triplicates. *p*<* 0.05 compared to untreated cells and ^#^p<0.05 with respect to infected cells (student’s *t*-tests paired analysis).

### Effect of glucose and mannose on 2-DG induced inhibition of virus multiplication

As shown in the previous section glycolysis is an intrinsic requirement of virus multiplication in host cells and its inhibition by 2-DG attenuated the virus growth. In a recent study, it was reported that enhanced glucose level favours SARS-CoV-2 growth [6]. To test the effect of increased glucose level on the growth inhibition efficacy of 2-DG, we treated the cells with additional 10 mM Glucose. The 3-fold increased glucose level (5.5 mM in media + 10 mM) enhanced the virus multiplication by 32% (Fig 6A) in infected cells. The 2-DG treatment (5 mM) inhibited the virus multiplication by 95% at equimolar glucose concentration in media, however, on the addition of 10 mM glucose this inhibition was significantly decreased to 88% as compared to its respective control and 84% with respect to infection control (Fig 6A). It is interesting to note that 3-fold increased glucose (∼15 mM) could reduce the effect of 2-DG (5 mM) by 7 %, only. This finding suggests that glucose is not the primary and efficient competitor of 2-DG uptake in virus-infected cells. Since 2-DG also shares structural similarity with mannose, an epimer of glucose, we tested if mannose can attenuate the effect of 2-DG on SARS-CoV-2 multiplication. Interestingly, we found that mannose is able to reduce the effect of 2-DG on virus multiplication at equimolar and 1/5^th^ of mannose (1 mM) to 2-DG (5 mM) ratio (Fig 6B). We further checked the effect of mannose on glucose uptake. In the qualitative and quantitative examination, it was found that mannose inhibited the uptake of 2-NBDG (a fluorescent analogue of glucose/2-DG) in infected cells (Fig. 6 C-D). These results suggest that the uptake of 2-DG is affected mainly by mannose and not much by glucose, which reduces the effect of 2-DG, on SARS-CoV-2 multiplication.

**Figure 6.**
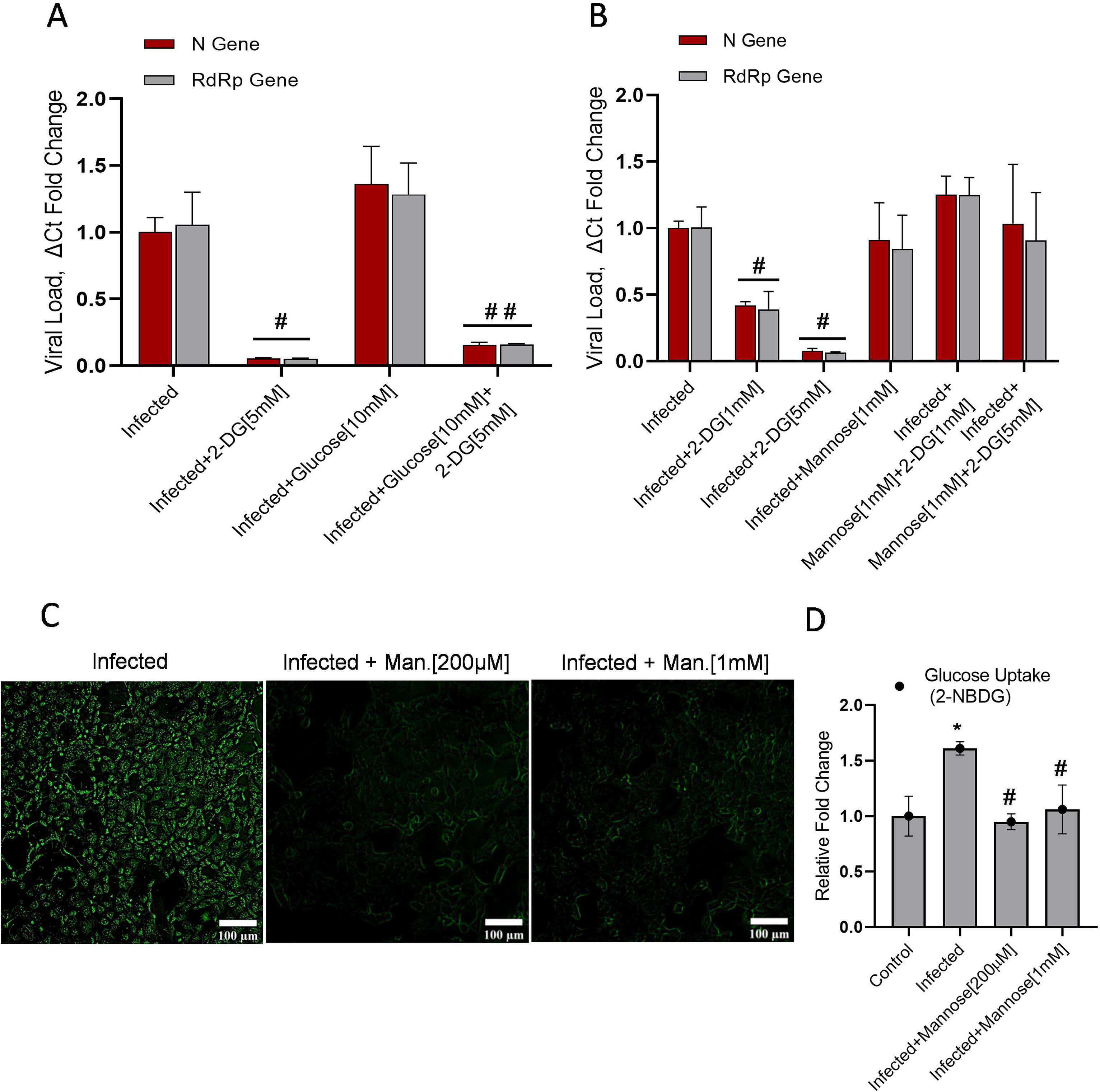
2-DG competes with mannose for intracellular uptake in SARS-CoV-2 infected cells A-B. The RFC graph presented with respect to the infected group alone indicating the 2-DG induced inhibition of Nuclear and RdRP gene and the magnitude of alteration in 2-DG efficacy showed by the separate combination of **(A)** glucose (glu) and **(B)** mannose (Man.) at the indicated concentrations. **C.** Photomicrographs shows the uptake of 2-NBDG [200µM] affected by equimolar concentration 200µM and 1mM of mannose at 48 h post-infection. **D.** In the similar experimental condition 2-NBDG uptake was measured on Spectrofluorometer using 465/540 (ex/em.) and the graph represents the RFC of 2-NBDG in the indicated groups. Data represent the mean ± SD of two independent experiments performed in triplicates. *p*<*0.05 compared to control; ^#^p<0.05 with respect to infected and ^##^p<0.05 compared to infected+2-DG (student’s *t*-tests paired analysis).

### Effect of 2-DG on Virus protein glycosylation and ER stress

Most envelop proteins of human pathogenic viruses are extensively glycosylated [5]. Various amino acids in the spike protein of SARS-CoV-2 was found extensively glycosylated and certain N-glycosylation sites of the receptor-binding domain (RBD) were found rich in both high mannose and complex type glycans [15, 16]. It is well evident from previous studies that 2-DG as an analogue of glucose and mannose interferes with protein glycosylation, resulting in misglycosylated or unglycosylated envelop proteins [5]. The mass spectrometry (MS) analysis of SARS-CoV-2 proteins from secreted virions revealed that in contrast to infected cells, the two N-glycosites (N331 and N343) of RBD are unglycosylated in 2-DG treated (at both 1 and 5 mM) cells following infection (Fig. 7A). Interestingly, highly mannosylated N331 and N343 glycosites of RBD are reported to be critical for the interaction of the virus at ACE2 receptor and entry to human cells [17]. Therefore, these results suggest that by interfering with N-glycan biosynthesis at crucial N-glycosites of RBD (N331 and N343), 2-DG may intervene in the SARS-CoV-2 binding at ACE-2 receptor and subsequent internalization of the virus at the host cell surface (Fig. 7B). Besides, 2-DG is known to induce Endoplasmic Reticulum stress (ER stress) by inhibiting glycolysis and glycosylation [18]. Given the observation on 2-DG induced unglycosylation of crucial residues in RBD, we analyzed if 2-DG treatment-induced ER stress in infected cells. Comparatively, a significant 1.4 to 1.6 fold increase in the levels of ER stress markers like GRP-78, ATF-6α, GADD-153 was observed in infected cells treated with 2-DG (Fig. 7C). The activation of p-eif-2α was also observed in the 2DG treated cells (Fig. 7C). These results indicate that 2-DG induced unglycosylation is causing ER stress in infected cells, which may interfere in virion packaging and production.

**Figure 7.**
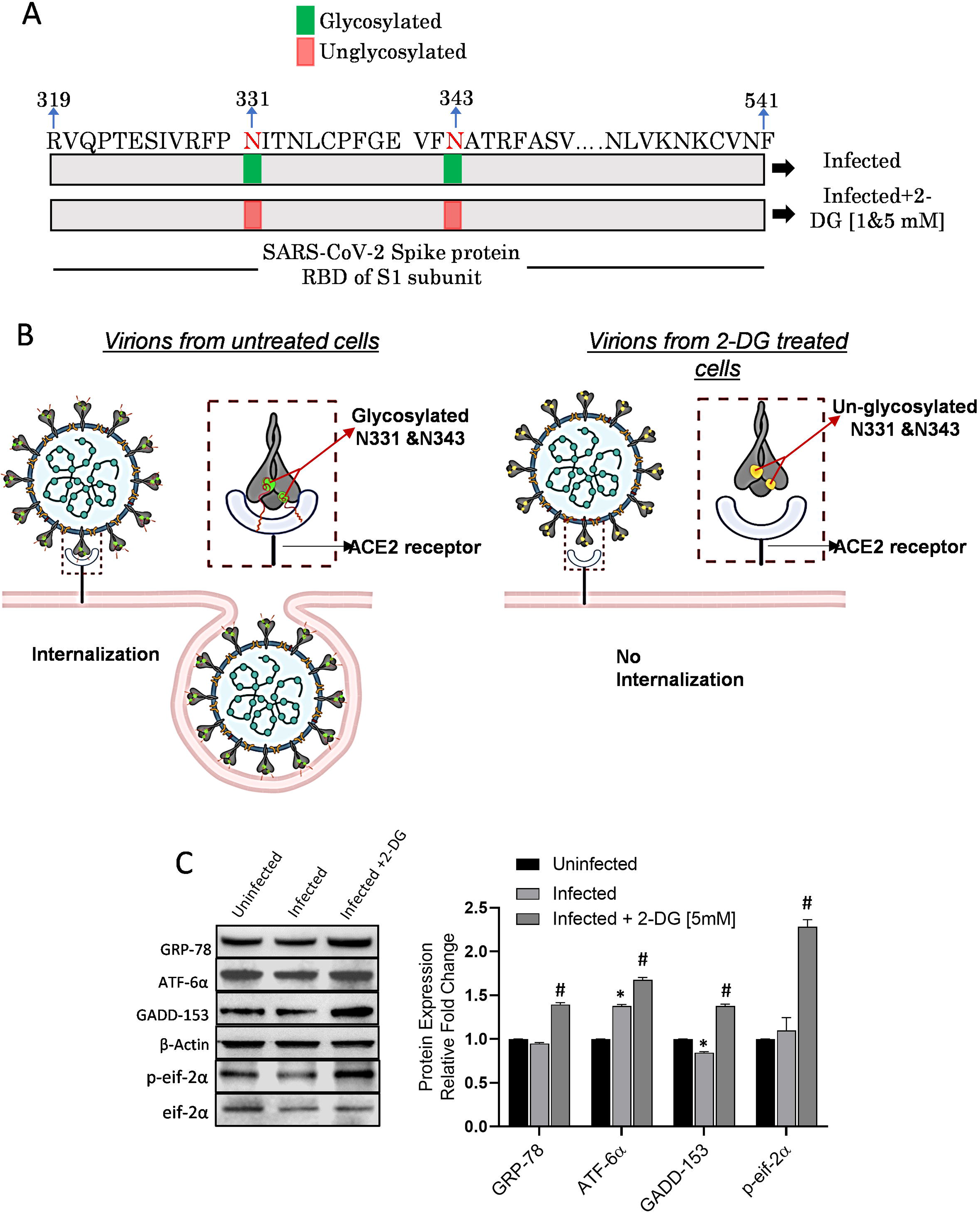
2-DG induces unglycosylation and ER stress. **A.** Schematic representation of N-glycosites (N331 and N343) on RBD (319-541) of SARS-CoV-2 S protein (S1 subunit) shows the glycosylation status in the 2-DG treated and untreated cells following SARS-CoV-2 infection. **B.** Graphical explanation of glycosylated and unglycosylated N-glycosites (N-331 & N-343) mediated virus-host interaction and internalization in untreated and 2-DG treated cells. **C.** Immunoblot and densitometric evaluation indicating the differential expression pattern of mentioned ER stress marker proteins in the untreated and 2-DG (5mM) treated cells following SARS-CoV-2 infection in Vero E6 cells. Data represent the mean ± SD of four independent experiments performed in triplicates. *p*<* 0.05 compared to uninfected and ^#^p<0.05 with respect to infected cells (student’s *t*-tests paired analysis).

### 2-DG weakens the infectivity potential of progeny virions

As per the published evidence, glycosylation of N331 and N343 glycosites of RBD is considered crucial for virus interaction and internalization to human cells [17]. Therefore we hypothesized that progeny virions produced from infected and 2-DG treated cells should have compromised infectivity. To test this hypothesis as an alternate mechanism of 2-DG for SARS-CoV-2 attenuation, we collected the secreted virions from media of untreated and 2-DG (1 & 5 mM) treated cells and measured their infectivity in fresh Vero E6 culture. After normalizing the viral concentration based on Ct values, we infected fresh cells with different virions samples (P3, refer to methodology section) without 2-DG treatment and analyzed the CPE and virus growth (Fig 8A). The progeny virions from 2-DG treated cells showed visibly reduced CPE at 48 hrs post-infection (Fig. 8A). This result was further confirmed by nearly 80% and 90% reduced virus growth estimated by RT-PCR in cells infected with progeny virions from 1 and 5 mM 2-DG treated cells, respectively (Fig. 8B). Additionally, these results were also substantiated by the protein expression analysis of N and S proteins in P3 progeny infected cell lysates from 2-DG treated (0.5, 1 & 5mM) and untreated cells. The cells infected with 2-DG treated progeny virions showed a dose-dependent decrease in the levels of N and S proteins indicating that virus protein synthesis is significantly compromised linked with the low infectivity of progeny virions compared to the untreated cells (Fig. 8C).

**Figure 8.**
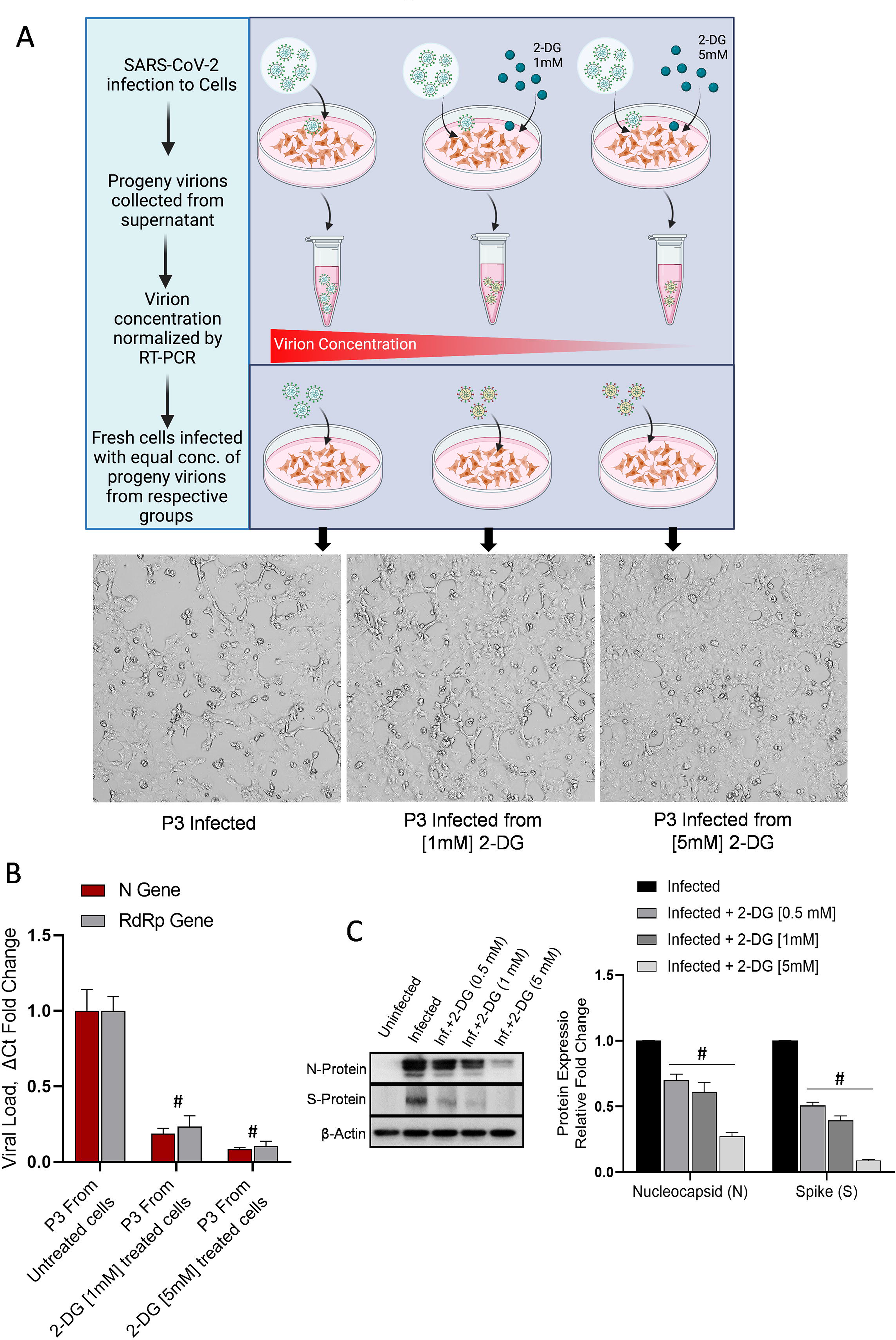
2-DG treatment produces defective progeny virions. The graphical depiction indicates the methodology followed for the experiment of progeny virion (P3) infection. The cells were infected with progeny virions and no further 2-DG treatment was given following infection. The photomicrographs show the progeny virions induced by CPE by bright field imaging **(A)** in the indicated treatment groups. **B.** The estimation of Nuclear and RdRP genes was performed in the P3 infected samples by RT-PCR assay. **C.** Immunoblotting of virus N and S protein was performed in cell lysate infected with P3 progeny virion of indicated treatment groups; The densitometry graph shows the RFC among indicated progeny fractions. **D.** Data represent the mean ± SD of four independent experiments performed in triplicates. ^#^p<0.05 compared to infected and untreated cells (student’s *t*-tests paired analysis).

## Discussion

Today whole world is struggling with the serious problem of the COVID-19 pandemic and looking for an effective therapeutic solution. The identification of mutations in the SARS-CoV-2 genome at a rapid rate and the possibility of mutations in prone genes is a huge concern targeting the efficacy of the vaccine as well as therapeutic resistance [7, 19]. Whenever new virus-induced pathogenesis is identified, it takes years to identify, characterize and develop a specific prophylactic and therapeutic approach against it. Therefore, to counter any virus outbreak at its commencement, there is a need to develop an effective broad-spectrum anti-viral strategy to prevent it from becoming an epidemic and/or pandemic.

Virus being an obligate parasite hijacks host cell machinery and reprograms it to favour for its own multiplication [5]. A compromised mitochondrial metabolism and an enhanced glycolytic pathway, upon SARS-CoV-2 infection, is one of the examples of virus infection-induced metabolic reprogramming [6, 7]. Normal/ uninfected cells show metabolic plasticity due to the presence of both glycolytic and mitochondrial metabolism. However, the glycolysis metabolic predominance in SARS-CoV-2 infected host cells makes it a lucrative target for developing therapeutic strategies. Therefore, we used a well known glycolytic inhibitor, 2-DG to counter the SARS-CoV-2 multiplication in host cells. This molecule has been tested extensively in lab and clinical trials.

We found that glucose uptake and ATP production was significantly upregulated in infected cells. This observation was also supported by enhanced protein levels of three major glucose transporters, GLUT1, GLUT3 and GLUT4 in infected cells (Fig. 1D). While GLUT1 and GLUT4 both have a medium-range affinity for glucose, GLUT3 exhibits a very high affinity for glucose with a very low Km value in the range of 1mM. Hence, GLUT3 plays a crucial role in the glucose transport of cell types with a high demand for energy in glucose hungry cells, even at a very low surrounding glucose concentration [20]. Therefore, in addition to the virus-induced levels of GLUT1/4, an upregulation of GLUT3 together ensure the high influx of glucose to infected cells even at the low concentration of extracellular glucose. In addition, increased levels of key glycolytic regulatory enzymes HK-II, PFK-1 and PKM-2 ensures the metabolism of glucose through glycolysis and other anabolic pathways. Further, an increased level of LDH-A confirms the Warburg phenotype of infected host cells (Fig. 1D). Since there is significantly higher glucose demand in SARS-CoV-2 infected cells, the intake of 2-DG (glucose mimic) will also be selectively high in infected cells at low 2-DG concentration, which was validated by increased uptake of 2-NBDG (at 0.2 mM, a fluorescent analogue of glucose/2-DG). 2-DG caused 30% to 40% inhibition of cell proliferation in the range of 1 to 5 mM with no significant effect on clonogenicity indicates that 2-DG is not cytotoxic/ genotoxic upto 5 mM (Fig 3A&B). Reduced 30% to 40% proliferation observed in 2-DG treated cells is due to the cytostatic effect of this molecule. Moreover, it is pertinent to note here that the inhibition of proliferation by 2- DG is not a matter of concern as the SARS-CoV-2 infected host cells in humans are primarily non-proliferating in nature and the cytostatic effect exerted by 2-DG is transient and can be reversed after removal of the drug [21].

Considering the selective higher uptake of 2-DG in SARS-CoV-2 infected cells and its potential to inhibit the virus multiplication, we tested its efficacy on B.6 variant at CCMB, Hyderabad in May 2020 and later on B.1.1 at INMAS, Delhi. 2-DG is able to significantly inhibit the virus multiplication at concentrations ranging from 0.5 mM to 5 mM, which was further verified by significantly reduced protein levels of SARS-CoV-2 nucleo-capsid protein (Fig. 4A-E) and reduced CPE (Fig. 5A). We observed profoundly high cytopathic effect and cell death in SARS-CoV-2 infected cells, which was significantly reversed by 2-DG treatment (Fig. 5A-C). Taken together these observations suggest that 2-DG inhibits the virus multiplication in infected host cells and thereby reduces the infection-induced CPE and cell death. However, 2- DG on its own did not contribute to cell death in this concentration range. This was further substantiated by the observation that 2-DG significantly reduced the ATP production in infected cells, however, it could not induce a meaningful reduction in ATP levels in uninfected cells (Fig. 4F). It is also pertinent to mention here that the cellular debris of dying cells is one of the major causes of inflammation, therefore reduced cell death in the 2-DG treated sample will result in reduced inflammation [22]. Supporting this, an earlier study on rhinovirus infection reported that 2-DG treatment reduces rhino virus-induced lung inflammation in a murine model [10].

A recently published study reported that enhanced glucose levels favour SARS-CoV-2 growth and thereby diabetes is turning to be one of the primary comorbid conditions responsible for the poor outcome in the treatment of COVID patients [6]. Therefore, we tested the effect of increased glucose on the efficacy of 2-DG. Irrespective of a 300% increase in glucose, the efficacy of 2-DG induced inhibition of virus multiplication was reduced by 7% (from 95% to 88%) at a given concentration of 2-DG (Fig. 6A). This result shows that the high relative glucose concentration could only minimally compromise the 2-DG efficacy, suggesting that the inhibition potential of 2-DG can be maintained at a low 2-DG to glucose ratio, also. The reversal of the inhibitory effect of virus multiplication by 2-DG and glucose uptake at an equimolar concentration of mannose (structurally similar to 2-DG), indicate that mannose can inhibit the 2- DG uptake and compromise the efficacy of 2-DG. It is noteworthy to mention here that the uptake of mannose can take place at very low plasma concentrations (30 to 50 micromolar) and cells use high-affinity glucose transporters like GLUT3 or dedicated mannose transporter which are insensitive to glucose [20, 23]. The important point to note here is that similar to mannose, 2- DG uptake can also take place even at its lower concentrations and a higher concentration of glucose may not be able to compromise the effectiveness of the 2-DG uptake and efficacy, profoundly. Our study provides a kind of evidence that 2-DG uptake is also mediated through either high-affinity glucose transporters like GLUT3 or selective mannose transporter in infected cells, which warrants further investigation.

N-glycosylation is a common feature of the viral envelope proteins [24]. SARS envelope proteins are reported to be highly glycosylated and responsible for the virus to host cell interaction, infection and immune evasion by glycan shielding. Being the obligate parasites, viruses are dependent on host-cell machinery to glycosylate their own proteins in the process of replication/multiplication [25]. In line with earlier observations on 2-DG induced de/mis- glycosylation of viral envelop proteins, we also found that two crucial residues N331 and N343 in RBD of the spike protein of the secreted SARS-CoV-2 virus from 2-DG treated cells were un- glycosylated (Fig. 7A). It is reported that glycosylation of N331 and N343 is critical for virus binding and infection in host cells [17]. This is further supported by the concentration-dependent reduction in infectivity of progeny virions collected from media of 2-DG treated cells (Fig. 8) suggest the defective and compromised infectivity potential of these virions, supporting the proposition that 2-DG induced un-glycosylation is leading to the production of defective/weaker virions probably resulting in reduced cell binding and internalization (Fig. 8). Altered glycosylation of proteins leads to accumulation of misfolded proteins in ER causing ER stress [18]. Increased level of ER stress markers (Fig. 7C) also indicates that 2-DG induced un- glycosylation may be leading to misfolding of viral proteins resulting in ER stress. Moreover, it was observed that the inhibition of viral multiplication by 2-DG was reversed upon addition of mannose (Fig. 6B), suggesting that mannose may inhibit 2-DG uptake (Fig. 6D) and reverse the glycosylation status of both glycosites (N331 and N343) leading to normal virus multiplication in host cells, even in the presence of 2-DG. These results strengthen the observation that 2-DG also inhibits the virus multiplication by interfering with glycosylation of spike protein of SARS-CoV-2. However, more information is required to establish the role of 2-DG induced un- glycosylation in the formation of defective virions. The doses at which 2-DG has been used in clinical trials, its maximum plasma concentration varies in the range of 1mM [26]. Therefore, it is also pertinent to note here that 1mM of 2-DG inhibits the virus multiplication by approximately 65% (0.35 fold remaining) and progeny virions produced from 1mM treated cells loses infectivity and result in 80% (0.2 fold remaining) reduced multiplication and secretion of the virus. Thus, the effective inhibition of SARS-CoV-2 multiplication obtained at 1mM 2-DG was 93% (0.35 x 0.2= 0.07 fold remaining).

## Conclusion

The findings presented in this manuscript highlights the SARS-CoV-2 infection mediated enhanced glucose metabolism in host cells, which can be targeted for therapeutic application. 2- DG exploits the inherent and natural mechanism of infected host cells for selective, high accumulation of the drug without compromising uninfected/ normal cell functioning, significantly. 2-DG by inhibiting both catabolic and anabolic pathways reduces the virus replication and infectivity of the progeny virions, which has a compromised potential of infection in neighbouring cells (Fig. 9). Although the effect of 2-DG has been analyzed on only 2 different SARS-CoV-2 variants (B.6 and B.1.1), its anti-viral property is suggested to be universal on all the variants of SARS-CoV-2, as 2-DG interferes with the metabolic requirement of virus-infected host cells. In summary, we demonstrate that glycolytic inhibitor 2-DG exhibits significant potential to be developed as a therapeutic to combat the COVID. These experimental evidences and previous clinical trial experience of 2-DG made way for this molecule to reach clinical trials in COVID-19 patients in India.

**Figure 9.**
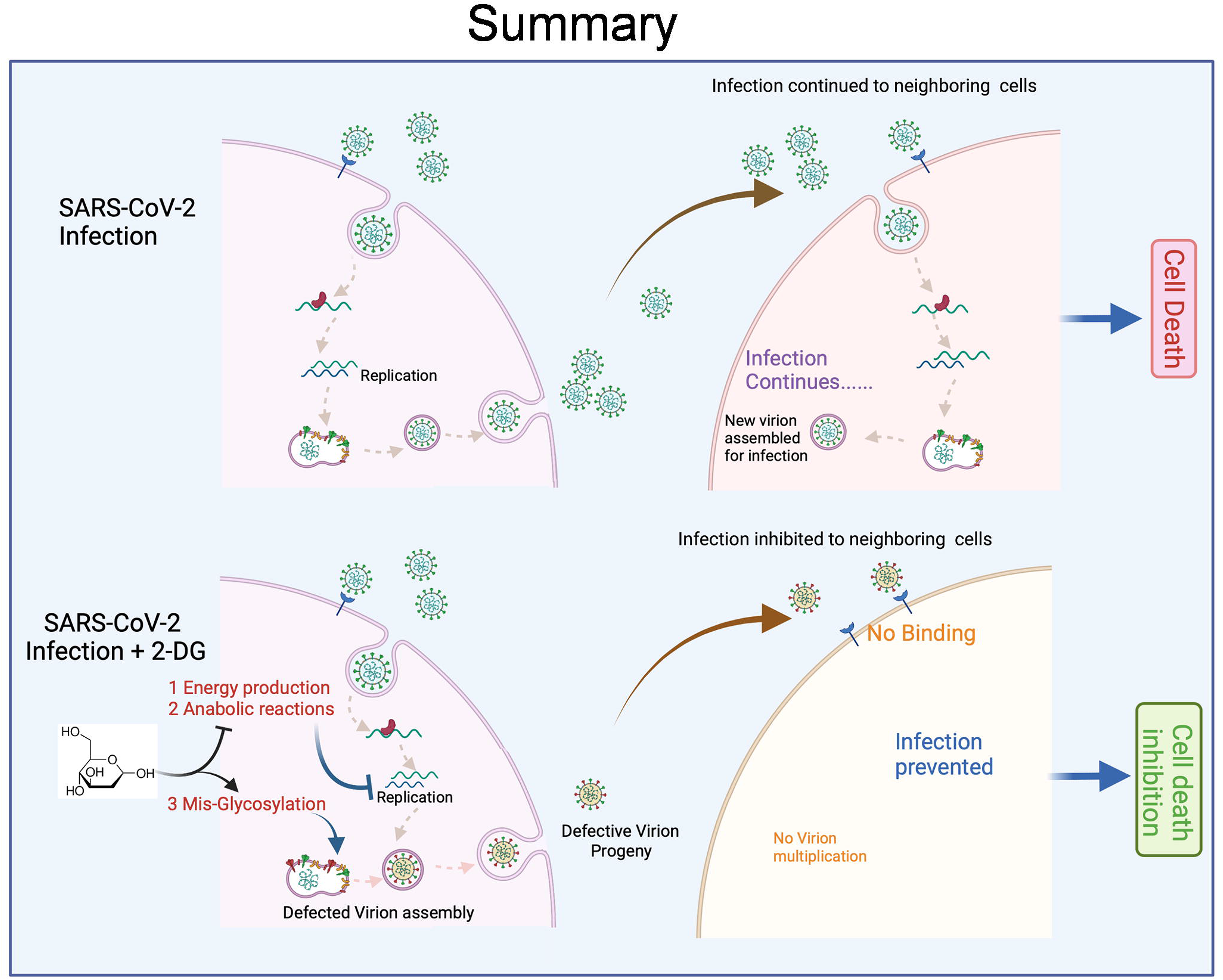
Mechanism of 2-DG induced inhibition of SARS-CoV-2 replication: The differential graphic representation of SARS-CoV-2 infection (upper half) and targeted inhibitory effect of anti-metabolite 2-DG (lower half) in the infected host cells.

## Material and Methods

### Cell culture, virus propagation and virus quantification

Vero E6 cell line was maintained at 37°C with 5% CO_2_ in an incubator, the culture was propagated in minimum essential medium (MEM; Sigma-Aldrich, St. Louis, MO) supplemented with 10 % heat-inactivated fetal bovine serum (FBS; Gibco, Life Technologies, Paisley, UK). The SARS-CoV-2 virus was isolated from a positively tested nasopharyngeal swab at BSL-3 facility INMAS-DRDO, Delhi. Briefly, Vero E6 cells were infected with filtered positive viral transport medium (filtered through 0.2µm filter) and mixed in the ratio 1:1 with MEM supplemented with 2% FBS, followed by incubation at 37°C in 5% CO_2_ with repeated mild agitation for 1h. The inoculum was removed post-incubation and the culture was washed with PBS and further, the resulting positive culture was re-supplemented with MEM having 10% FBS and maintained at 37°C & 5% CO_2_ up until cytopathic effects were apparent in the cells. The resulting SARS-CoV-2 viral stock (P1) was then collected and confirmed by RT-PCR. The viral stock was used for one more passage to obtain a working stock (P2) which was then used for all the experiments. Viral titrations for P2 was performed in Vero E6 cells seeded in 6-well plates at a density of 0.75x10^6^ cells/well using plaque enumeration as previously described with some small modification [21]. Plaque forming units (PFU) were calculated using the formula: PFU/ml = the number of plaques/ (Dilution factor * Volume of diluted virus/well). The multiplicity of infection (MOI) was derived from the formula: MOI = PFU of virus used for infection/ No. of cells and in the present study the MOI was kept low (0.3), which was standardized to ensure the experimental study and optimal analysis of SARS-CoV-2 induced intracellular stress behaviour in Vero E6 cells. A similar protocol was followed for establishing B.6 variant in Vero cells, at the BSL-3 facility, CSIR-CCMB, Hyderabad.

### Virus growth kinetics and Bright-field imaging

For SARS-CoV-2 infection, cells were seeded into 24 well-plates with the density of 0.05×10^6^ cells/well and were infected with MOI of 0.3. For extracellular virus RNA quantification, media was collected from each well at 24 h and virus RNA was isolated using MagMAX Virus/Pathogen II Nucleic Acid Isolation Kit (Thermo Fisher Scientific) and automated extraction was carried out using KingFisher™ Flex Purification System, KingFisher with 96 Deep-well Head (Thermo Fisher Scientific). For intracellular virus RNA quantification, TRIzol^®^ Reagent (Ambion, Life Technologies) was used, further isolation and extraction were performed as mentioned earlier. Multiplex rRT-PCR kit (Allplex^TM^ 2019-nCoV assay, Seegene) based identification of the three target genes associated with SARS CoV-2, i.e. Nuclear (N) gene, RNA dependent RNA polymerase (RdRP) gene and Envelop (E) gene were performed following the manufacturer’s protocol in CFX96 touch real-time PCR detection system (Biorad). Moreover, morphological changes associated with cytopathic effects (CPE) caused by SARS-CoV-2 infection in Vero E6 cells were observed at 24h and 48h post-infection under Bright Field (10X objective) on an inverted microscope (Nikon ECLIPSE Ts2).

### Analysis of infectivity

The comparative analysis of infection potential of progeny virions (P3) from SARS-CoV-2 infected untreated and 2-DG treated cells was examined using RT-PCR estimation of N and RdRp genes from the supernatant of respective samples of Vero E6 cells. The P3 virions in the supernatant were quantified by RT-PCR from infected untreated and 2-DG treated cells and obtained values were used to normalize the concentration of virions in each group. Then freshly cultured Vero E6 cells were infected with a normalized equal concentration of virions (P3) from each group. Further N and RdRP gene analysis was performed to examine the infectivity potential of P3 virions.

### Estimation of glucose uptake and ATP determination

The qualitative and quantitative estimation of glucose uptake was performed using 2-NBDG (200µM) in Vero E6 cells. Similarly, for differential glucose uptake in Vero E6/Vero cells initially, Vero cells (0.025×10^6^) were stained with CellTracker^TM^ Red CMPTX (Invitrogen) using (5µM; 30mins; 37°C) followed by normal growth medium replacement and co-culture with unstained Vero E6 cells (0.025×10^6^). Briefly, cells were seeded with density 0.05×10^6^ cells/well in 24 well plate. The next day infection was carried as described in the previous section and 2-DG, glucose (glu.) and mannose (Man.) treatment as mentioned in the respective figure legend. Glucose uptake was monitored at 48 h post-infection by incubating cells with 2- NBDG (200µM; 20mins; 37°C) probe buffer solution containing MgCl_2_ and CaCl_2_ (1mM each) in PBS. Further cells were washed twice with PBS and fluorescence images were captured under a fluorescence microscope (Nikon ECLIPSE Ts2-FL) using 10×10 X magnification. The quantitative estimation of 2-NBDG fluorescence was carried out at 465/540 (ex/em.) and ATP production was measured at 560nm Luminescence on Synergy H1 Hybrid Multi-Mode Microplate Reader (BioTek Instruments, USA) on Synergy H1 Hybrid Multi-Mode Microplate Reader (BioTek Instruments, USA) followed by respective group cell number normalization. Subsequently, all the fluorescence images were subjected to morphometric analysis to quantify the intracellular uptake of the fluorophore. For this purpose, the study employed algorithms from CELLSEGM and Image Processing toolbox in Matlab 2020a (Mathworks). It initially segmented the foreground (nucleus, cell) from background pixel intensity and segmented more than 1000 cell objects from multiple fields of view and the segmented cell object’s median pixel intensity was calculated [27].

### Cellular proliferation efficiency and clonogenicity

Sulforhodamine B assay was employed to derive relative proliferation efficiency of Vero E6 cells upon administration of 2-Deoxy-D-Glucose (2-DG). Vero E6 cells were seeded in a 96 well plate at a density of 0.001x10^6^ cells/well and were left overnight in a CO_2_ incubator and were treated with concentrations of 2-DG ranging from 0.1mM to 100mM and the assay was performed as previously described method [28]. CFSE (Sigma) staining for cellular proliferation was performed in accordance with the manufacturer’s protocol. Vero E6 cells stained with CFSE (5µM; 20 min, at room temp.) in MEM with 2% FBS. After incubation, cells were washed with a normal growth medium and reseeded 35mm petri dishes (PD) with the density of 0.25x10^6^ Cells/PD and kept overnight in dark in a CO_2_ incubator. 2-DG treatments were given the following day, and the cells were terminated and processed for flow cytometry (BD LSR II) based analysis at 0h, 24h and 48h. Macrocolony assay was performed for determining the plating efficiency of 2-DG treated cells at 1mM and 5mM in Vero E6 cells. Cells were seeded in 60mm petri dishes and were treated with 2-DG for 24 h. Further, these cells were reseeded with 200 cells/PD with the control group and kept for 7-8 days in a humidified incubator to allow the formation of macroscopic colonies. After that cell were washed once with PBS and stained with crystal violet followed by an enumeration of colonies consisting of >50 cells. Finally, the plating efficiency was calculated as the ratio of (Number of colonies formed/ Number of cell-seeded)×100 and presented as % clonogenicity.

### Apoptosis assay

Apoptosis assay was performed using ethidium bromide (EB)/acridine orange (AO) staining in Vero E6 cell line at 48 h post-infection in the treatment groups (as mentioned in the respective figure legend). Vero E6 cells were seeded in 24 well plate (0.05×10^6^cells/well) and kept at 37°C in a CO_2_ incubator. The next day cells were infected with SARS-CoV-2 and 2-DG treatment was carried out (as described in the previous section). At the required time point cells were dislodged by trypsinization and floater were collected followed by centrifugation (100g; 10mins) and each group’s cells were transferred to a 96 well plate. Further cells were stained with AO/EB dye in the ratio of 1:1 (100µg/ml each) to each well. Finally, cell images were captured using a fluorescence microscope (Nikon ECLIPSE Ts2-FL) using 10×10 X (objective and eyepiece) magnification. Finally, the acquired fluorescence images of AO/EB cells were subjected to the morphometric analysis to estimate the median area of AO/EB uptake in fluorescence images of uninfected, 2-DG treated, infected, and infected +2-DG treated cells. For this purpose, the study employed the algorithms from CELLSEGM and Image Processing toolbox in Matlab 2020a (Mathworks). At first, fluorescence images were smoothened and subjected to the segmentation process to isolate cells having little overlap with other cells. This cell segmentation was carried out at many fields of view images, and more than 1000 non-overlapping cell objects were isolated, and their median area was calculated for each uninfected, 2-DG treated, infected, and infected with 2-DG treated category.

### Immunoblotting

Immunoblotting was performed to examine the differential expression of glucose transporters, glycolytic proteins and drug-induced change in virus nucleocapsid proteins as mentioned in the respective figure legend. Briefly, cells were seeded at 0.5×10^6^ density in 60mm tissue culture dishes and kept in a CO_2_ incubator at 37°C. The following day, virus and drug treatments were performed (as described previously). At the time of analysis whole cell lysate was processed in RIPA buffer containing Tris/HCl: (50 mM; pH 7.4), Na_3_VO_4_ (1 mM), EDTA (1 mM), NaCl (150 mM), NaF (1 mM) PMSF (2 mM), NP-40 1%; supplemented with protease inhibitor cocktail (1X), and protein was estimated by BCA method. Equal quantities of lysates (40µg and 60 µg was used for detection of nucleocapsid protein and glycolytic proteins respectively) were resolved on 10% or 12% SDS-PAGE gel (depending on the molecular weight of the respective proteins). Further protein samples were electro-blotted onto PVDF membrane (MDI) followed by membrane blocking with 5% BSA for 1 h (according to the manufacturer protocol). Incubation of the primary antibody was performed in the dilution of (1:1000; according to the manufacturer instructions) for 14 h at 4°C then washed in Tris-buffered saline supplemented with 0.1% Tween-20 (TBST) followed by HRP conjugated secondary antibody incubation (1:2500) for 2 h. Finally, blots were washed again with TBST and developed using ECL chemiluminescence detection reagent on Luminescent image analyzer (Image Quant LAS 500, Japan). Densitometry was performed for each blot using Image J software and obtained values were normalized with respective loading control β-actin, and the graph presented as relative fold change among the treatment groups.

### Virus harvesting, inactivation and glycosylation study

Vero E6 cells were infected with SARS-CoV-2 as mentioned in the previous section followed by 2-DG treatment (1 and 5mM). Virus harvesting, purification and inactivation were performed using the experimental protocol as described elsewhere [29] with some modifications. Briefly, the cell culture supernatant (25 ml) of infected and 2-DG treated cells were collected at 72 hrs post-infection and stored at -80°C overnight followed by centrifugation (4500 g; 10mins.). Further supernatant was overlayed on 20% sucrose (w/v), HEPES (25mM), in PBS and ultracentrifugation was performed at 103,745 g for 5hrs using Beckman Coulter Euro Center SA. Finally, the virus pellet was dissolved in TNE buffer (10 mM Tris, 0.2 M NaCl, 10 mM EDTA in PBS; pH 7.4) and inactivated by incubating samples at 56°C for 10 min. in Tris–HCL (100 mM) supplemented with SDS (5%). For the glycosylation study, proteolytic digestion of 30µg protein was performed and the obtained peptides were separated using liquid chromatography and eluates were directly infused into Electrospray ionization mass spectroscopy setup followed by the acquisition of MS and MS/MS data. Raw data was mapped with the SARS-CoV-2 (S) protein FASTA sequence and the analysis of peptide mapping and glycosylation status was carried out using BioPharma Finder software.

### Statistical analysis

All experiments were conducted at least three times unless otherwise indicated. Analysis of statistical significance between two groups was performed using Student’s t-test (paired analysis) and differences were considered significant when value p was < 0.05.

## Acknowledgements

We are thankful to Dr. Rakesh Mishra, Director CSIR-CCMB for facilitating the 2-DG testing against SARS-CoV-2 at BSL-3 facility of CCMB during nation wide lockdown. We are thankful to Dr Sankar Bhattacharya from THSTI and Dr Chandru from RCB, Faridabad for their help in providing Vero and Vero E6 cell line for virus culture. We are also thankful to Dr Jubilee Purkaystha, in-charge of BSL-3 facility INMAS for helping us in conducting the experiments at this facility. We also want to express our gratitude to Dr Viney Jain and Dr B S Dwarakanath for useful discussions and advice. AK is a recipient of a fellowship from CSIR.

