## Supplementary figures and images for "Glycolytic inhibitor 2-Deoxy-D-glucose attenuates SARS-CoV-2 multiplication in host cells and weakens the infective potential of progeny virions"

### Supplementary figure

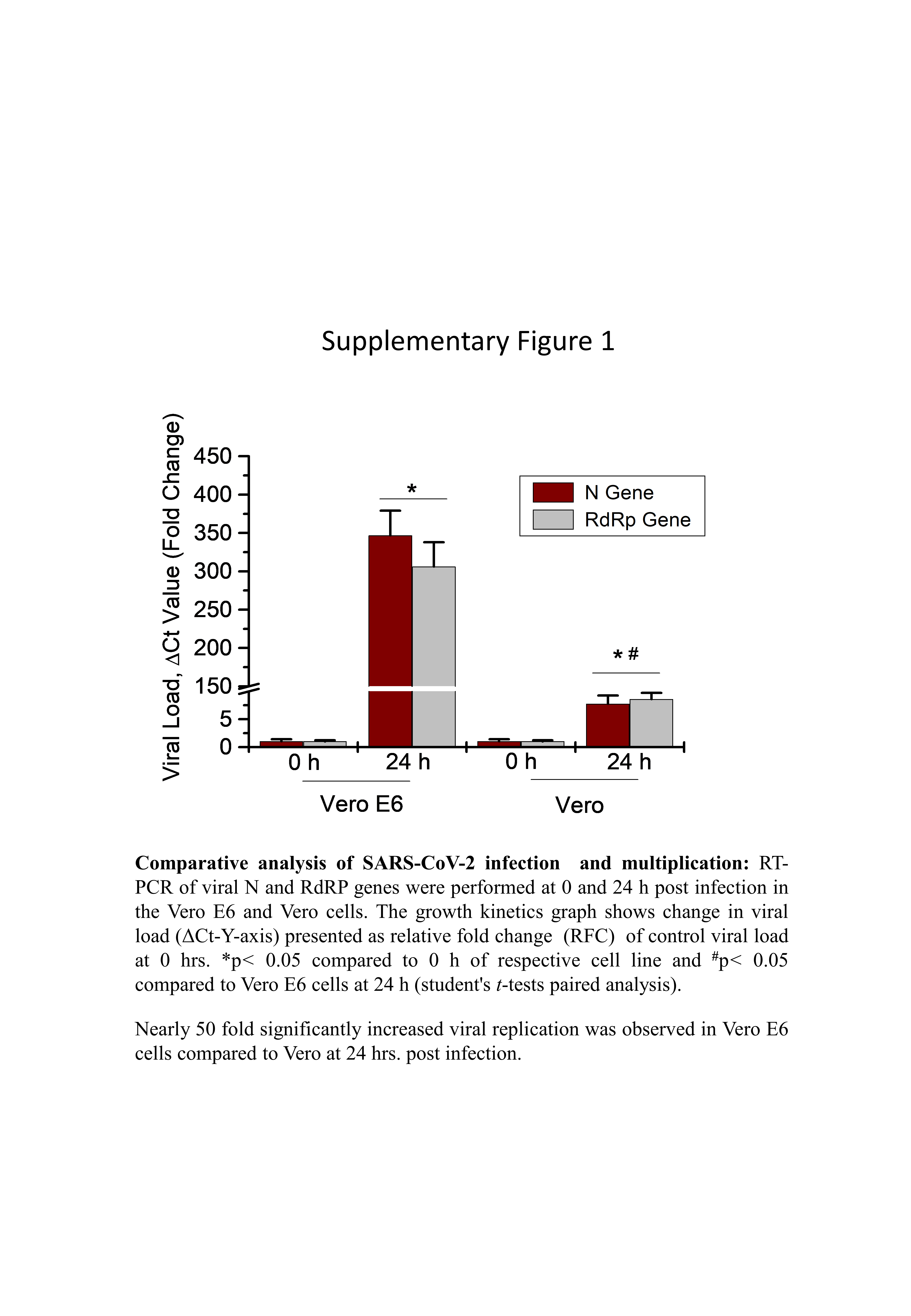
